# Plant parasitic cyst nematodes respond to viral infection through RNA interference

**DOI:** 10.64898/2025.12.02.691819

**Authors:** Hein Overmars, Matthijs Mobach, Stefan J.S. van de Ruitenbeek, Jaap-Jan Willig, Joris J.M. van Steenbrugge, Aska Goverse, Mark G. Sterken, Lisa van Sluijs, André Bertran

## Abstract

Plant-parasitic nematodes (PPN) cause yield losses to multiple important crop species and counteractive measures are only partially successful. Recently the first viruses that infect PPN were discovered, presenting new research opportunities to understand and halt PPN. A critical step in this process is to identify how PPN respond to viruses. Here, two PPN species, the potato cyst nematode *Globodera rostochiensis* and the beet cyst nematode *Heterodera schachtii,* were used to study PPN-virus interactions. Viral discovery based on bioinformatic analyses indicated three putative viral genomes present in *G. rostochiensis* (a Toti-like, Xinzhou-like, and Picorna-like virus) and five putative viral genomes in *H. schachtii* (a Bunya-like, Tobamo-like, Picorna-like, and two Nyami-like viruses). A genome composition analysis and RT-PCR data confirmed that these viruses infect the nematodes and not their host plants. Subsequently, antiviral RNAi activity was detected in both *H. schachtii* and *G. rostochiensis* using small RNA sequencing. For *G. rostochiensis* two distinct genotypes were investigated. These strongly differed in their RNAi response: Ro5-line 22 produced roughly 40 times more antiviral viRNAs than Ro1-line 19. Our measurement of an active antiviral response represents the first indication of antiviral activity in PPN.

**Author summary:** Plant-parasitic nematodes (PPN) are key agricultural pests and although some PPN viruses have been discovered recently, it is still unknown if and how PPN respond to viral infection. Here we find three viruses in the potato cyst nematode (*Globodera rostochiensis*) and five viruses in the beet cyst nematode (*Heterodera schachtii*). Of these viruses, six are newly discovered. Genome and molecular analyses confirm that all found viruses specifically infect nematodes and not their host plants. By measuring small RNA profiles, it was confirmed that PPN react to viral infection by means of RNAi, except for a weak response against two negatively stranded Nyami-like viruses. Moreover, distinct RNAi profiles were found for two potato cyst nematode lines, giving the first insight into which genes might be involved in RNAi in PPN. Together, this research provides the first proof of an active antiviral response in PPN. Thereby this finding opens new opportunities for biological control and gene editing tools.

## Introduction

Plant parasitic nematodes (PPN) are microscopic roundworms that specialize in feeding on plants and cause severe yield loss across multiple crop species worldwide [1,2]. Amongst these, cyst nematodes of the genera *Globodera* and *Heterodera* belong to the most harmful PPN for agriculture [1]. *Globodera* and *Heterodera* nematodes migrate intracellularly through the roots, damaging the plant root systems, before they form a feeding site. Withdrawal of nutrients by the nematode then leads to diminished plant growth due to malnutrition and deformations in the root architecture [3]. On top of that, these nematodes produce many offspring which survive encapsulated in the dead female body (the cyst) in the soil for years. Diverse management options exist, although these all have substantial disadvantages such as environmental harm (nematicides), limited durability or application (resistance genes), workload (crop rotation, soil hygiene measures), or efficacy (biocontrol). Accumulating studies showing that PPN can be infected by viruses encourages new research directions [4–10], for example in the field of biocontrol or creation of transgenic tools. However, for working towards such applications it is imperative to know if these viruses evoke an antiviral response in their hosts.

It was not before 2011 that the first virus infecting a nematode was confirmed, namely, the Orsay virus that infects the free-living nematode *Caenorhabditis elegans* [11]. From there on, thanks to rapid progress in next generation sequencing (NGS) technologies, multiple reports on the identification of viruses in different PPN species were published. At least eight different viruses were found in PPN by obtaining sequences of the potato cyst nematodes *Globodera pallida* and *Globodera rostochiensis*, and the soybean cyst nematode *Heterodera glycines* [4–7]. Then viruses infecting the beet cyst nematode *Heterodera schachtii* and the root lesion nematode *Pratylenchus penetrans* were identified [8,10]. Lastly, a study on viral discovery in nematode communities extracted from agricultural soils in the USA was performed [9]. Independent wet-lab evidence of these viruses has mostly been generated by RT-PCR. Next, for some of the PPN viruses, localization studies via fluorescence-based or classical *in-situ* hybridization methods were conducted [7,8]. Many of the discovered viruses appear throughout the nematode’s life, from egg to adult, suggesting these viruses are vertically transmitted within nematode populations [4,6,12]. But how the PPN respond to the presence of viral infection remains unknown.

So far, molecular nematode-virus interactions have been studied in *Caenorhabditis* nematodes, especially the *C. elegans*-Orsay virus patho-system. Within *C. elegans*, three antiviral pathways respond to viral infection. First, terminal uridylation can target the 3’ end of viruses leading to degradation [13]. The second pathway is called the Intracellular Pathogen Response (IPR), that not only responds to viruses, but also other intracellular parasites such as microsporidia and oomycetes [14–17]. This genetic pathway is expanded in *C. elegans* and, so far, appears to be restricted to *Caenorhabditis* spp. [14,18]. The third antiviral pathway is called RNA interference (RNAi) and targets exogenous RNAs, including those of viral origin, for degradation [11,19,20]. The antiviral activity of the RNAi pathway presents conserved virus-associated small interfering RNA (viRNA) patterns in *Caenorhabditis* spp., namely the accumulation of host-amplified antisense viRNAs of 22nt in length with a heavy bias towards a G nucleotide as the first base in the viRNAs [20–22]. Contrary to the IPR pathway, RNAi is found broadly across eukaryotic organisms, including PPN. PPN including *Heterodera* spp. and *Globodera* spp. are capable of RNAi as indicated by RNA soaking, host-induced gene silencing and comparative genomics [23–28]. Thus, it is likely that if the PPN viruses provoke a host response, antiviral RNAi will be part of it.

In this study, the molecular antiviral response was investigated in *Globodera rostochiensis* and *Heterodera schachtii*. Initially, eight near-complete viral genomes were discovered, including six novel viruses. Three viruses were discovered in two *G. rostochiensis* isogenic lines (Ro1-line 19 and Ro5-line 22) [29] and five viruses were discovered in the *H. schachtii* Woensdrecht isolate [30,31]. We were able to confirm the presence of these viruses in nematode populations by RT-PCR, and to clone and sequence their genomes. To ascertain the host species of these viruses, genome composition analyses (CpG/UpA) and phylogenetic analyses were conducted. These results indicated these viruses specifically infect nematodes and are not plant viruses transmitted by nematodes. Finally, we sequenced the small RNAs produced by infected nematodes. Hereby we demonstrate for the first time that PPN produce an active antiviral response against the viral infection.

## Materials and methods

### Nematode populations

For *Heterodera schachtii* (Beet Cyst Nematode; BCN) a Woensdrecht isolate (kindly provided by Stichting IRS, Dinteloord, the Netherlands) was used. This Woensdrecht isolate of BCN was maintained in the greenhouse in a continuous culture on white cabbage; *Brassica oleracea* (cultivar “Cyrus”). For *Globodera rostochiensis* (Potato Cyst Nematode; PCN) four populations/lines were used; Ro1-Mierenbos, Ro5-Harmerz and their inbred lines Ro1-line 19 and Ro5-line 22, respectively. These lines were selected by *in vitro* controlled single matings [32]. Cysts were mass multiplied in de greenhouse on susceptible potato cultivar “Eigenheimer” and long term stored at -80°C [32]. Cysts from these PCN populations were transferred from - 80°C to 4°C at least 2-3 months prior to use [33].

### Harvesting of different life stages

Pre-parasitic second stage juveniles (J2s) of PCN were hatched by incubating cysts in tap water for 7 days followed by incubation in root exudate from *Solanum lycopersicum* cv. Money Maker for 3 days. For BCN, J2s were obtained by incubation of cysts in 3 mM ZnCl_2_ for five days. To obtain eggs from BCN, cysts were gently crushed in a mortar and pestle. The freed eggs were sieved through a 100 µm (pore diameter) sieve. Eggs and hatched Juveniles (J2s) of BCN as well as J2 of PCN were sucrose purified, washed extensively in tap water and transferred to 1.5 mL RNase free tubes.

Second generation females of BCN were collected from a greenhouse culture on *Brassica oleracea* cabbage plants (cultivar “Cyrus”). Cabbage plants were grown in pots and inoculated with hatched J2s. After 74 days, females were collected as follows: root balls were gently cleaned from soil with tap water and further cut into 2-3 cm pieces and transferred onto a stack of sieves (top to bottom); 1000, 365 and 180 µm pore diameter. Females were rinsed from the roots with a strong beam of tap water and gravid females collected on the 365 µm sieve while young females were collected on the 180 µm sieve. Females were hand-picked and collected in a 1.5 mL RNase free tube. All nematode samples were incubated in RNA-later (Invitrogen) for 30-45 minutes at room temperature (ca. 20°C). Next, most of the RNA-later was removed by pipetting and samples were stored at -80°C.

As a control, roots of non-infected 3 weeks old *Brassica oleracea* (cultivar “Cyrus”) plants were washed until free of soil particles, surface dried between paper towel and snap frozen in liquid nitrogen. Roots of 6 plants were pooled and ground in liquid nitrogen until a fine powder. The root powder was aliquoted in 1.5 mL RNase free tubes and stored at -80°C.

### RNA extraction

Nematode and root material was homogenized as follows: To each sample 2 steel beads (diameter of 3 mm) and 35 µL of homogenisation buffer (Maxwell LEV16 Plant RNA kit, Promega) were added and homogenized in a tissue homogenizer (Qiagen GmbH, Swingmühle Tissue Lyser 2) at 30/s for 2x 80 seconds. Then, 100 µL of homogenisation buffer (Maxwell) was added and samples were again homogenized at 30/s once for 80 seconds. The steel beads were removed immediately after homogenization. Subsequently, RNA extraction was performed according to the Maxwell LEV16 plant RNA kit (Promega) protocol as provided by the manufacturer. Later, RNA concentration was measured in a Qubit 3 Fluorometer with the Qubit™ RNA High Sensitivity (HS) Assay Kit, according to the manufacturer’s instructions and RNA integrity was analysed on a 1% agarose-TAE gel. Only RNA samples that showed a 2:1 ratio for the ribosomal RNA genes 28S and 18S respectively were used in this study.

### RNAseq datasets used and or generated for viral discovery and small RNA analysis

Virus discovery in BCN was performed on the RNAseq dataset of Willig *et al*., 2022 (samples 19-29; ERR9604282 – ERR9604293 except ERR9604283) and in PCN on the RNAseq samples from *G. rostochiensis* Ro1-line 19 and Ro5-line 22, (SRR16693885 and PRJEB66473, respectively) (**Table 1**) [29,30]. The BCN RNAseq dataset was generated from poly-A enriched total RNA and the PCN RNAseq dataset were generated through ribosomal RNA depletion.

**Table 1.**
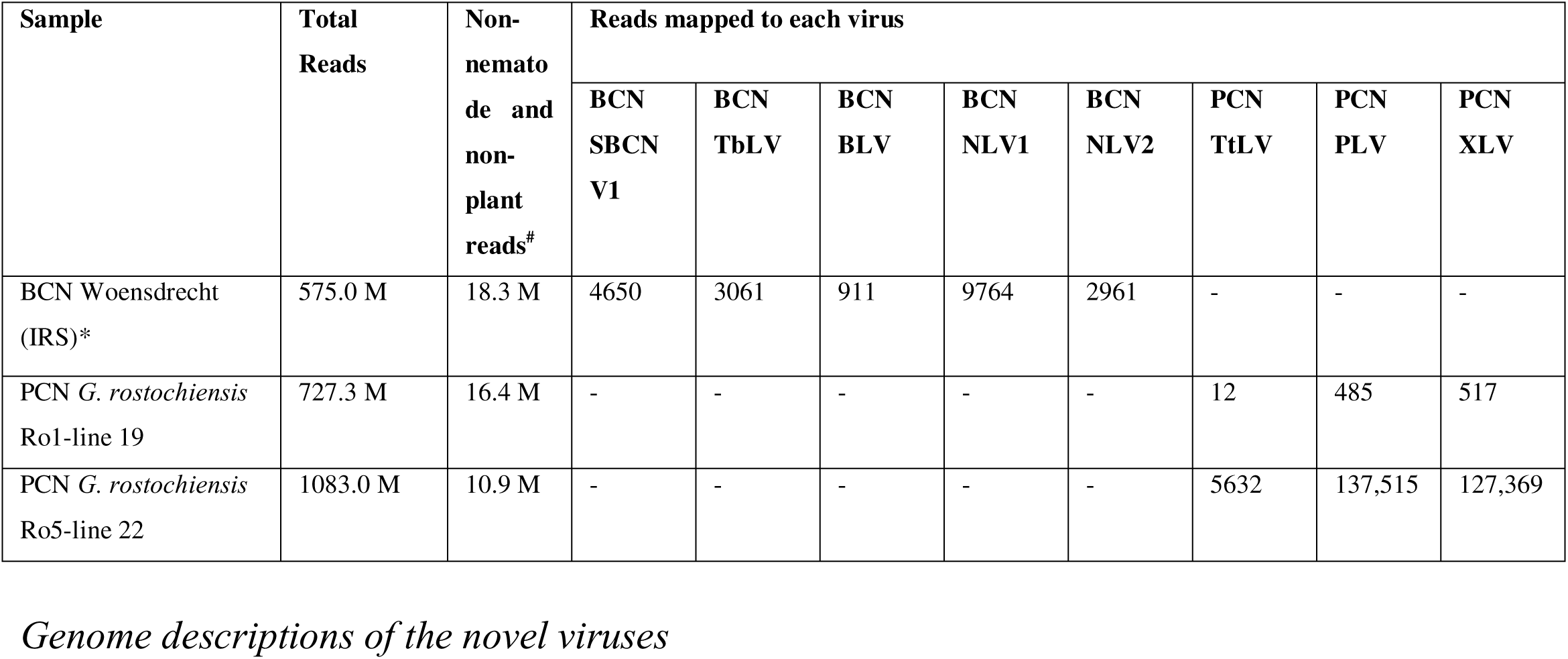
Viral reads detected in *G. rostochiensis* and *H. schachtii*. Number of reads derived from Illumina RNAseq reads mapped to each virus discovered in BCN Woensdrecht and PCN *G. rostochiensis* Ro1-line 19 and Ro5-line 22 populations. *For BCN Woensdrecht a total of 10 samples were pooled together as this dataset consists of total RNA extracted from BCN parasitizing *Arabidopsis* plants. The PCN samples were obtained from pre-parasitic J2s.

For small RNA-enriched RNAseq libraries total RNA from *G. rostochiensis* Ro1-line 19 and Ro5-line 22 and *H. schachtii* Woensdrecht were used for size selection, library preparation (NEBNext small RNA library - NEB) and RNA sequencing (Illumina HiSeq 2500 SE 50) by Macrogen and were deposited under PRJEB66168.

### RNAseq virus discovery pipeline

Paired-end Illumina reads for all BCN and PCN samples were mapped using *hisat2* version 2.2.1 [34] and *samtools* version 1.14 [35] against the genomes of *H. schachtii* and *G. rostochiensis* lines Ro1-line 19 and Ro5-line 22 [29,31] as well as the genomes of their host plant *A. thaliana* (in the case of the BCN [30]) while using snakemake version 8.25.4 [36]. The RNAseq datasets of *G. rostochiensis* were generated from hatched J2s directly from our cyst collection. Reads that did not map to the host plant or nematode genomes were kept after mapping using command ‘samtools view -c -f 4’, transformed from bam to fastq using bedtools *bamtofastq* version 2.30.0 [37]. The fastq files of the BCN and PCN’s were fed into SPAdes for the *de novo* transcript assembly using the ViralRNASPAdes algoritm of SPAdes version 3.15.2 [38] with the command ‘--rnaviral’ within spades.py. *De novo* assembled transcripts were then used as query for local BLASTn/x version 2.11.0 [39] against a collection of all Ecdysozoan viruses obtained from NCBI Virus (March 2021). Additionally, the top 10 longest-assembled transcripts were manually analysed for the presence of ORFs and conserved viral protein domains using CLC Main Workbench (version 23). BLASTn and BLASTp [39] were also performed for the top 10 transcripts. Incomplete or partially *de novo* assembled genomes were extended with the contig assembly tool of CLC Main Workbench (v. 23) by matching these sequences with the bulk of the *de novo* assembled transcripts generated by SPAdes from the corresponding dataset. Lastly, the pool of non-host reads was mapped against the near complete viral genomes using hisat 2.2.1 with the help of samtools utilities using snakemake. Mapped reads were visualized with IGV [40] and counted using samtools idxstats 1.21.1 [35].

### DNAseq analysis for viral genome integration

Long and short DNA reads generated for *G. rostochiensis* Ro1-line 19 and Ro5-line 22 and BCN Woensdrecht were used as queries to look for viral presence in nematode genomic DNA raw reads [29,31]. Mapping of the viral genomes to long genomic DNA reads was performed using minimap2 v. 2.1 [41] and samtools 1.14 [35] and to short genomic DNA reads was performed using bwa-mem2 version 2.2.1 [42] with the help of samtools utilities using snakemake version 8.25.4 [36]. The final alignments were visualized with IGV version 2.14.1 [43].

### Small RNAseq analysis

Small RNA samples were generated from the same total RNA used for the amplification of conserved viral RdRp domains, determination of viral UTRs (explained below) and RNAseq by size selection enrichment (performed by Macrogen using the NEBNext small RNA library prep kit). Small RNA Illumina reads were first trimmed to remove adapter sequences using CutAdapt. Reads where then filtered by size and only those between 15 and 35 nt in length were kept. These were then mapped to the near full-length viral genomes generated by our virus discovery pipeline (see above). Reads were mapped using the BWA and mapped reads were separated into forward and reverse mapped and further filtered with the tag XM < 2 (i.e., less than two mismatches in the whole read) using SAMtools view and Filter BAM in local Galaxy v.22.01. Pileup outputs of the aligned reads over each genome were generated using the bcftools plugin of Galaxy with all settings on. All data processing was performed in Local Galaxy [44] while graphs were generated in RStudio 2023.06.1 using the *Tidyverse* package [45].

### Amplification of conserved viral RdRp domains and determination of viral UTRs

Between 800-900 ng (Qubit) of Total RNA from artificially hatched J2s, eggs, young- and gravid females, and cabbage roots was used for cDNA synthesis using either random hexamer or oligo-dT (12-18) primers with GoScript (Promega) according to the manufacturer’s instruction. PCR reactions were performed in 25 µL reaction volume with 0.02 U/µL Phusion™ High-Fidelity DNA Polymerase (Thermo Fisher Scientific), 1x Phusion HF buffer, 0.2 mM dNTP mix and 0.5 µM of each forward and reverse primer (**Supplementary Table S1**) targeting the conserved RdRp domain of the respective viruses. PCR conditions in all reactions were: denaturing at 98°C for 30 seconds followed by 35 cycles of 98°C for 10 seconds, 61°C for 20 seconds and 72°C for between 40 seconds – 100 seconds, followed by a final extension for 3 minutes at 72°C. RT-PCR end products were analysed on a 1% Agarose-TAE gel using a 1Kb+ DNA ladder (Invitrogen) for estimating product sizes. Fragments were excised from the gel, cloned into pCR™ 2.1-TOPO™ TA vector (Invitrogen) and sent for Sanger sequencing (EZ Sequencing, Macrogen) according to the manufacturer’s protocols. Sequences were analysed and alignments made with sequence analyser program BioEdit (v7.2.5).

The 5’ and 3’ ends of the viral genomes of BCN TbLV, BCN NLV1, BCN NLV2, PCN PLV and PCN XLV were determined using the SMARTer RACE 5’/3’ Kit (Takara Bio USA) using total RNA (1 µg) extracted as described above. In short, 5’-first strand cDNA synthesis was performed using random primers. 3’-first strand DNA synthesis required an additional step because the newly discovered viruses may lack a polyadenylated (polyA) tail. Therefore before 3’-first strand cDNA synthesis, polyA tails were added to the RNA using the Poly(A) Polymerase (Takara Bio USA) at 37°C for 60 min followed by a 5-minute termination step at 85°C. A clean-up step using NucleoSpin columns (Macherey-Nagel, Germany) was essential for follow-up reactions to generate 3’-first strand cDNA. Then, 5’- and 3’-first strand cDNA was used for Rapid Amplification of cDNA ends (RACE) using a touchdown PCR followed by a secondary ‘nested’ PCR according to the manufacturer’s instructions (primer sequences in **Supplementary Table S1**). Finally, products were gel extracted, cloned into a provided pRACE vector and Sanger sequenced (EZ Sequencing, Macrogen) according to the manufacturer’s protocols.

### Phylogenetic analysis

The open reading frames (ORFs) identified in the novel viral genomes were determined using CLC Main Workbench v.23. The ORFs corresponding to the RdRp gene of each virus or the viral polyprotein, when applicable, where translated into amino acid sequences and submitted to NCBI Virus. Deposited viral genomes sharing similarities to the novel viral genomes were obtained from this repository and used for multiple protein alignments generated with MUSCLE in CLC Main Workbench v.23. Maximum Likelihood trees were calculated using the web servers of IQ-TREE [46] with best model selection including AIC and BIC corrections. Free-rate heterogeneities were allowed, and an ultrafast bootstrap of 10000 was used. Trees were edited in iTOL [47]. Only branches with a bootstrap score above 70 were kept.

### Gene orthology analysis of the RNAi pathway genes

The complete proteomes of our three populations of cyst nematodes were investigated for the presence of orthologues of the genes in the RNAi pathway as known from *C. elegans*. To this end, Orthofinder was used to find orthogroups for each of the proteins in the RNAi pathway [48]. As additional species/populations we used the complete proteomes of *Ditylenchus destructor* (PRJNA312427), *Bursaphelenchus xylophilus* (PRJEB40022), *Meloidogyne chitwoodii* (PRJNA666745), *Heterodera schachtii* Bonn (PRJNA722882), *Heterodera glycines* (PRJNA381081), *C. remanei* (PRJNA577507), *Trichinella spiralis* (PRJNA12603) and *O. volvulus* (PRJEB513). All proteomes were obtained from WormBase Parasite v. WBPS18. In parallel, local BLASTp (from the BLAST+ package) was run for all the proteins of *C. elegans* against our three nematode populations.

### Genome composition analysis

Genomes of viruses infecting free-living nematodes, plant-parasitic nematodes and plants were downloaded from NCBI (for NCBI reference numbers, see **Supplementary Table S3, S4**). Fragments were checked for sequencing quality by the percentage of unassigned (N) nucleotides. All had less than 0.1% unassigned nucleotides and were used in subsequent analyses.

Genomes compositions were extracted using the package ‘Biostrings’ [49] and custom written functions in R (v4.2.1) were made to calculate the frequency of nucleotide (*Freq_i_*) and dinucleotide (*Freq_ij_*) pairs in the genome according to

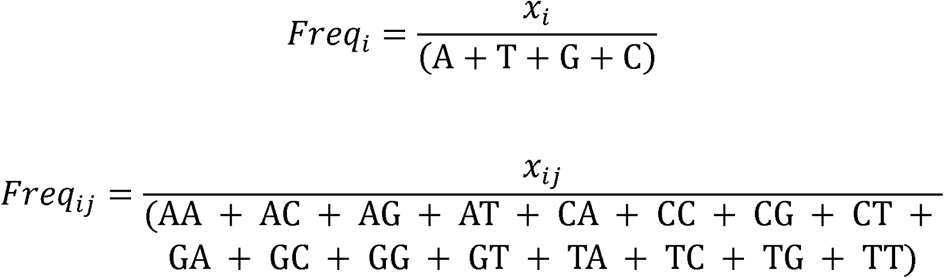

where x_i_ is a specific nucleotide or nucleotide combination (A, T, C, G) and x_ij_ is a specific dinucleotide combination (AA, AC, AG, AT, CA, CC, CG, CT, GA, GC, GG, GT, TA, TC, TG, TT).

Dinucleotide observed-to-expected (O/E) ratios (R_O/E_) were calculated by

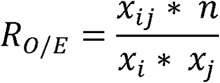

where x_ij_ is the dinucleotide pair (e.g. CG), x_i_ is the first nucleotide (e.g. C) of the dinucleotide pair and x_j_ is the second dinucleotide (e.g. G) of the nucleotide pair and n is the sequence length.

A principal component analysis (PCA) was performed on nucleotide, dinucleotide and observed-to-expected dinucleotide ratios using the *prcomp* function with normalized variance (scale. = TRUE) of the ‘Stats’ package.

## Data availability

The small RNAseq datasets generated in this work have been deposited in Biostudies under the reference number PRJEB66168 (run ERR12050910, ERR12050911, and ERR12050912). The genome sequences of BCN BLV, BCN TbLV, BCN NLV1, BCN NLV2 and SBCNV1 found in BCN Woensdrecht and PCN PLV, PCN XLV and PCN TtLV are deposited in Genbank (PQ140468, PQ140469, PQ140470, PQ140471, PQ140472, PQ140473, PQ140474, PQ140475). Data analyses performed in R can be found on: https://git.wur.nl/published_papers/overmars_sluijs_2024_ppn_viruses.

## Results

### Discovery of novel beet cyst and potato cyst nematode viruses

Driven by curiosity, we identified putative viral sequences in our laboratory populations of the beet cyst nematode (BCN) *Heterodera schachtii* (BCN; Woensdrecht isolate) and inbred populations of potato cyst nematode (PCN) *Globodera rostochiensis*: Ro1-line 19 and Ro5-line 22 [32]. We chose to investigate these nematode species because they belong to the most studied PPN species, were previously shown to carry viruses [12,50], and because of available reference genome assemblies [29]. A virus discovery analysis was performed on published RNAseq data for *H. schachtii* and on two RNAseq data sets generated from J2s of *G. rostochiensis* Ro1-line 19 and Ro5-line 22 (SRA database references SRR16693885 and ERP15124, respectively) [30,31]. These analyses identified five putative near full-length viral genomes in *H. schachtii* Woensdrecht and three putative near full-length viral genomes in *G. rostochiensis* Ro5-line 22, while only few and comparatively short viral fragments were obtained from *G. rostochiensis* Ro1-line 19 (**Table 1**). None of the viruses discovered in *H. schachtii* could be observed in *G. rostochiensis* and vice versa.

### Genome descriptions of the novel viruses

To predict the structures of the putative viral genomes we performed multiple BLAST alignments and Maximum-Likelihood-based phylogenetic analyses for either their polyprotein or polymerase genes (**Figure 1, Supplementary Figure 1**). Moreover, in order to generate complete viral genomes, 5’and 3’ RACE were performed to obtain and possibly extend the untranslated regions (UTRs) obtained via *de novo* assembly (**Supplementary Table S2**). Our results indicate that the *de novo* assemblies yielded high quality UTRs which were only marginally extended by the RACE analyses. Additionally, it was investigated if any of the near full-length viral sequences were present in the genome of the nematodes. We found no genomic DNA reads mapping to any of the viral sequences we identified. This indicates that these viral genomes most likely represent autonomous viruses instead of endogenized viruses or viral elements.

**Figure 1.**
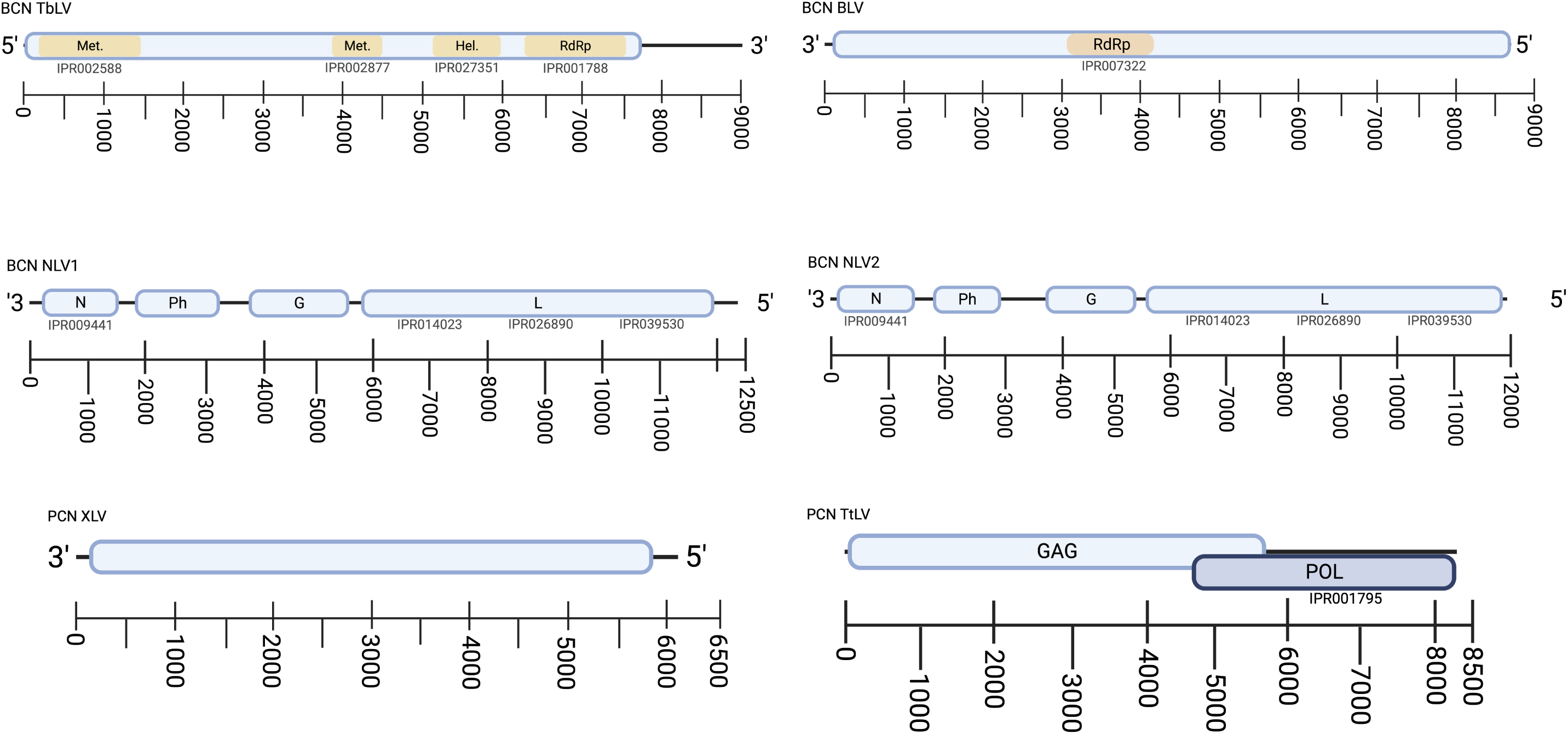
Genome structures of the novel viruses identified in BCN Woensdrecht and PCN Ro5-line 22 based on their near full-length structures. From the five viruses identified in BCN Woensdrecht, one is near identical to SBCNV1 [10] and was therefore not presented here as we do not consider it a novel virus. For PCN Ro5-line 22 the picorna-like virus PCN PLV is 89% identical in amino acid sequence to PCN PLV described by Ruark *et al*., 2018 [12] and for this reason it is not shown here as a novel virus. Viral genomes and their open reading frames are displayed according to their actual size in the sequences. BCN NLV1, BCN NLV2, BCN BLV and PCN XLV are all negative single-stranded RNA viruses and their (non-coding) viral (v)RNA is presented here from 3’ to 5’. This strand is used by the viral polymerases for viral mRNA transcription. Genes expressed from these mRNAs are aligned/superimposed on the negative strand vRNA shown. BCN TbLV is a positive single-stranded RNA virus and PCN TtLV is a dsRNA virus, and both are shown in the (coding) 5’ or 3’ end orientation. The line below each of the genome structures indicates the size of the viral genomes in nucleotides. Conserved protein domains are highlighted either by brown inlays in the case of a large ORF (open reading frames) and indicated by their Interpro codes right below the protein names otherwise.

First, the five viruses discovered in BCN were considered and are hereafter described as a Picorna-like virus (SBCNV1), two Nyami-like viruses (BCN NLV1 and BCN NLV2), a Bunya-like virus (BCN BLV) and a Tobamo-like virus (BCN TbLV). The Picorna-like virus discovered shares 98% protein identity with the previously discovered sugar beet cyst nematode virus 1 (SBCNV1, YP_009553470.1) and is therefore considered to be a new isolate of the same species [10]. The two novel Nyami-like viruses (BCN NLV1 and BCN NLV2) contain all functional domains typical for Nyami-like viruses (**Figure 1**), except for the open reading frame coding for protein III (M, matrix protein) that is present in the Nyami-like virus discovered in soybean cyst nematodes (SCN-NLV) [12]. Phylogenetically BCN NLV1 clusters closely with the partial sequence of the SCN-NLV virus isolate (AVK42876.1) found in beet cyst nematodes [12] (**Supplementary Figure 1**). Because the ∼1.8kb SCN-NLV fragment has 99.8% overlap in nucleotides with the ∼12.5kb BCN NLV1, this previously discovered SCN-NLV isolate may be a partial sequence. BCN NLV2 clusters with the three SCN-NLV isolates previously discovered in soybean cyst nematodes (AVK42865.1, AVK42870.1, AVK42875.1), but shares only 81% of protein identity, hence separating BCN NLV2 as a new species. BCN NLV1 and BCN NLV2 share just 71% protein identity. BCN BLV has a genome coding for a single large RNA-dependent RNA polymerase (L), and contrary to the soybean cyst nematode bunya-like virus SBCNV2 [10] we could not identify reads resembling an S or M RNA genome for BCN BLV. Moreover, the L protein of BCN only had 27% protein identity with SBCNV2 (KY609505) [12], hence we suggest BCN BLV is a novel virus. For BCN TbLV we identified conserved Interpro protein (functional) domains for most of the viral proteins typical for Tobamo-like viruses [51]. Tobamo-like viruses belong to a virus family for which no other virus species were previously described to be associated with nematodes. BCN TbLV is phylogenetically more closely related to a crustacean virus, Beihai charybdis crab virus 1, and to plant associated Tobamo-like virus 1 [52], rather than to other nematode-associated viruses discovered (**Supplemental Figure 1**) [9].

Based on found similarities, the three putative viral genomes (**Figure 1**) in *G. rostochiensis* Ro5-line 22 were identified as a Picorna-like virus (PCN PLV), a Xinzhou nematode virus 3-like virus (PCN XLV) and a Toti-like virus (PCN TtLV). From these three viral genomes, only PCN PLV seemed related to a previously described virus, PCN PLV (accessions MG550272 from *Globodera rostochienis* and MG550273 and MG550274 from *Globodera pallida*), which were 80-89% identical in nucleotide sequence to our new found PCN PLV [12]. In our phylogenetic analyses PCN PLV was found to cluster closely to these other PCN PLV isolates, albeit in a sister clade (**Supplementary Figure 1**). This phylogenetic placement indicates that the PCN PLV we found might carry sufficient genetic differences to be a different type of picorna-like virus infecting PCN. Notably, our phylogenetic analysis places all PPN-associated picorna-like viruses known so far in a very close relationship, contrary to picorna-like viruses infecting free-living or animal parasitic nematodes [8,12,53]. For PCN XLV we did not find conserved protein domains, and it represents a novel viral genome closely related to two previously discovered Xinzhou nematode virus 3-like viruses [9]. These viruses were the only nematode-associated viruses in a group containing insect-infecting Qin or Qin-like viruses (**Supplementary Figure 1**). The genome of PCN-TtLV, consists of two partially overlapping open reading frames identified as GAG and POL that encode for the capsid protein and the viral polymerase, respectively. PCN TtLV phylogenetic placement revealed that it is the sole nematode-associated virus in a clade containing multiple arthropod (ticks, insects, crustaceans) toti-like viruses. Hubei toti-like virus 5, a virus associated to a Spirurid parasitic nematode is also represented in this phylogeny albeit on a more distant relationship to PCN TtLV (**Supplementary Figure 1**) [54].

### Molecular verification of the novel PPN viral genomes

To experimentally verify the presence of the five *H. schachtii* viral genomes total RNA was extracted from different life stages of *H. schachtii* grown in white cabbage (*Brassica oleracea*) plants. The conserved RdRP domains predicted in each of the viral genomes identified were amplified and cloned and sequenced for further confirmation. Amplicons of the expected sizes were present in eggs, hatched J2, young females (> 180 µm and <385 µm) and fully developed females (> 385µm) while no amplicons were detected in the cabbage roots where these nematodes were propagated (**Figure 2A**). The presence of the five *H. schachtii* associated viruses in eggs, J2s and adult females, suggests vertical transmission of these viruses.

**Figure 2.**
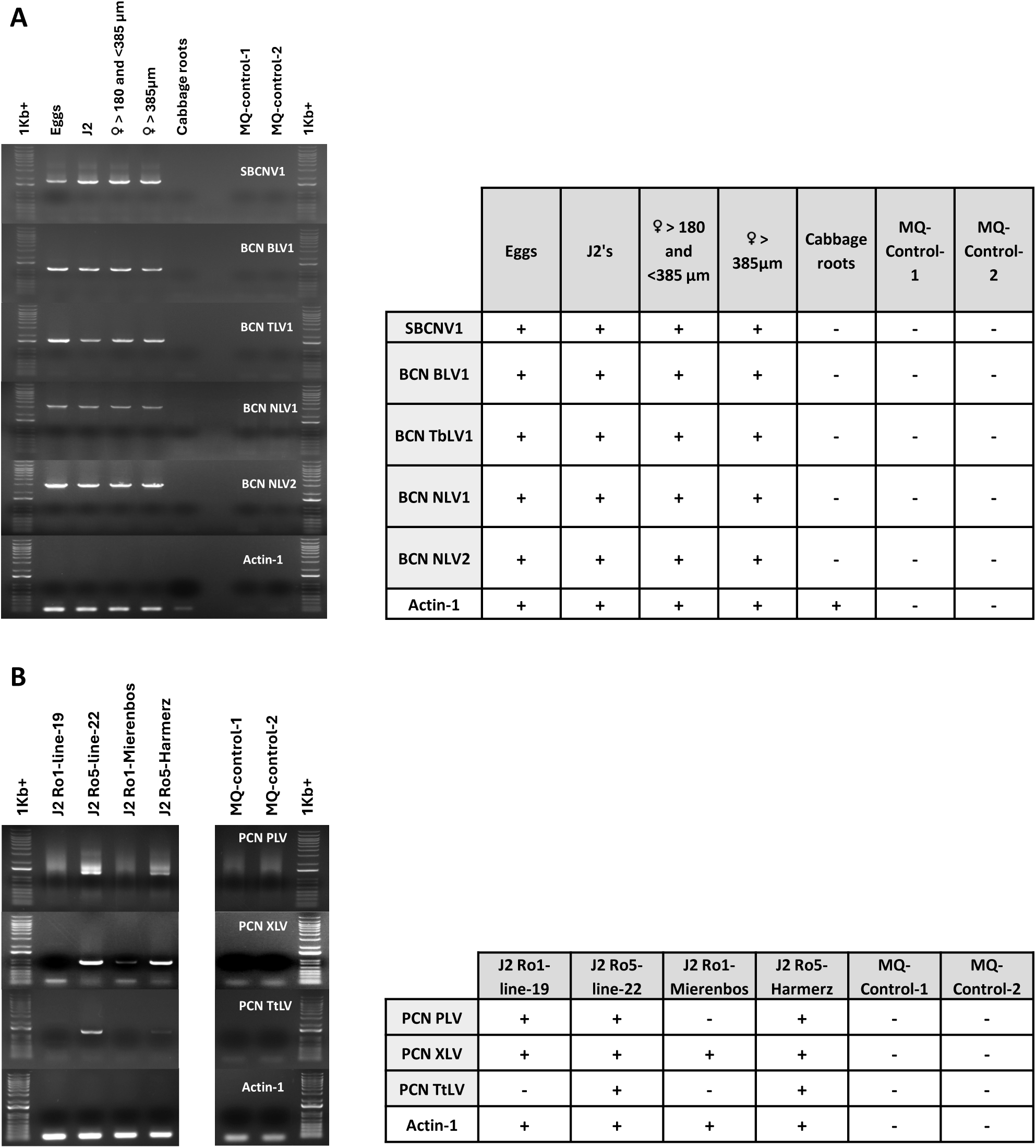
Confirmation of novel viral genome detections in BCN (A) and PCN (B) by RT-PCR. **A**) Detection of viruses in different lifecycle stages of BCN (*H. schachtii*, Woensdrecht isolate) by endpoint RT-PCR. Primers targeted the conserved domain of the RdRp of respective viruses. Expected PCR product sizes are: BCN NLV1 – 3120 bp, BCN NLV2 – 2812 bp, BCN TbLV1 – 1541 bp, BCN BLV1 – 1092 bp, SCNV1 - 1742 bp. Nematode Actin 1 gene specific primers (Gen-Act-F4 and Gen-Act-R4; 139 bp, target both Actin-1 of BCN and cabbage root) served as positive control in each RT-PCR test. MQ controls served as negative, non-template, controls. Uninfected cabbage root was included to test any of the viruses to be present in the nematode’s host. **B**) Detection of viruses in different lines of PCN. Ro1-line 19 and Ro5-line 22 are inbred lines derived from Ro1-Mierenbos and Ro5-Harmerz respectively. Primers targeted the conserved domain of the RdRp of respective viruses. Expected PCR product sizes are: PCN PLV - 1283 bp, PCN XLV – 925 bp, PCN TtLV1 – 1263. Nematode Actin 1 gene specific primers (Gen-Act-F4 and Gen-Act-R4; 139 bp target Actin-1 gene of PCN) serve as positive control in each RT-PCR test. MQ controls served as negative, non-template, controls. For clarity next to the panel the presence or absence of PCR end products is represented as + and - respectively, because of the poor visibility of some of the PCR products in the gel.

Confirmation of the three viral genomes identified in *G. rostochiensis* was performed on the J2 lifestage of Ro1-line 19, Ro5-line 22 and their parental populations, Ro1-Mierenbos and Ro5-Harmerz, respectively (**Figure 2B**). The conserved regions of the RdRPs of the three viral genomes were amplified in Ro5-line 22 and in its parental population, Ro5-Harmerz. PCN PLV and PCN XLV were amplified in Ro5-line 22, Ro5-Harmerz and in Ro1-line 19, despite there being no near full-length viral genome assembled from the latter population in our in *silico* analysis (**Table 1**). While the presence of PCN PLV could be confirmed in Ro1-line 19, it could not be confirmed in its parental population Ro1-Mierenbos. PCN TtLV was found neither in Ro1-line 19, nor its parental population, Ro1-Mierenbos. Finally, marked differences in virus accumulation were seen in the RT-PCRs performed on Ro5-line 22 and Ro1-line 19: Ro5-line 22 has much stronger amplification of the viral fragments than Ro1-line 19 (**Figure 2B**). The difference in the number of viral reads between these lines matches with the expectations, because fewer reads were found using the virus discovery pipeline in Ro1-line 19 (**Table 1**, **Figure 2B**).

### Genome composition analyses supports that the newly discovered viruses infect nematodes rather than plants

Nematode-associated viruses can infect the nematode itself, but can also use a nematode as a vector to reach new host species [55,56]. During the genome structure analysis, we noted that none of the open reading frames of any of the viruses we identified had a conserved movement protein. A movement protein would be expected if these were plant-infecting viruses vectored by nematodes [57]. Hence, we hypothesise that these autonomous viruses infect nematodes rather than plants. To strengthen our hypothesis, we performed a viral genome composition analysis. In this analysis we aimed to distinguish nematode-infecting viruses from plant-infecting viruses (vectored by nematodes or through other means of transmission) by comparing the genomic frequencies of observed/expected CpG and UpA dinucleotides. Plant-infecting viruses mimic the reduced CpG observed-to-expected (O/E) ratio of the plant mRNAs, likely to prevent being recognized as a virus by the host cell machinery [58,59]. Contrary, invertebrates such as nematodes do not reduce mRNA levels in their genomes and viruses infecting invertebrates contain CpG_O/E_ ratios in line with the frequency of C and G nucleotides in their hosts genome (CpG_O/E_ ratio ≈ 1) [59]. Additionally, UpA ratios are often similar between hosts and virus. Here, the genome composition of the newly obtained viruses was assessed using principal component analysis and by calculating CpG_O/E_ and UpA_O/E_ ratios to predict if these represent *bona fide* nematode-infecting viruses.

Publicly available genomes of viruses infecting free-living nematodes (*Caenorhabditis* spp.), viruses infecting plant-parasitic nematodes (*Heterodera* spp.*, Globodera* spp., and *Pratylenchus penetrans*), plant-infecting viruses using nematodes as vectors (*Xiphinema* spp.*, Longidorus* spp.) and the newly discovered viruses were analysed using principal component analysis (PCA) (**Figure 3A, Supplementary Table S3, S4**). The PCA was based on (di)nucleotide frequencies and O/E ratios. Additionally, viruses belonging to the order *Bunyavirales* (in particular from the former family of *Bunyaviridae)* were indicated as these often show aberrant genome characteristics as they can often infect both vertebrates and invertebrates [59]. As expected, members of the *Bunyavirales* cluster together on the PCA. The plant-infecting viruses cluster closely together, suggesting that these contain specific genome characteristics. Contrary, the viruses infecting free-living nematodes (FLN), plant-parasitic nematodes (PPN), and most of the newly discovered viruses from this study scatter throughout the plot. Notably, four viruses cluster with the plant-infecting viruses. These viruses belong to the family of *Picornaviridae*: both the previously described PCN PLV [12], SBCNV1 [7] and their strains identified in this study. In addition, the BCN NLV1 (*Nyamiviridae*) also clusters with the plant-infecting viruses.

**Figure 3.**
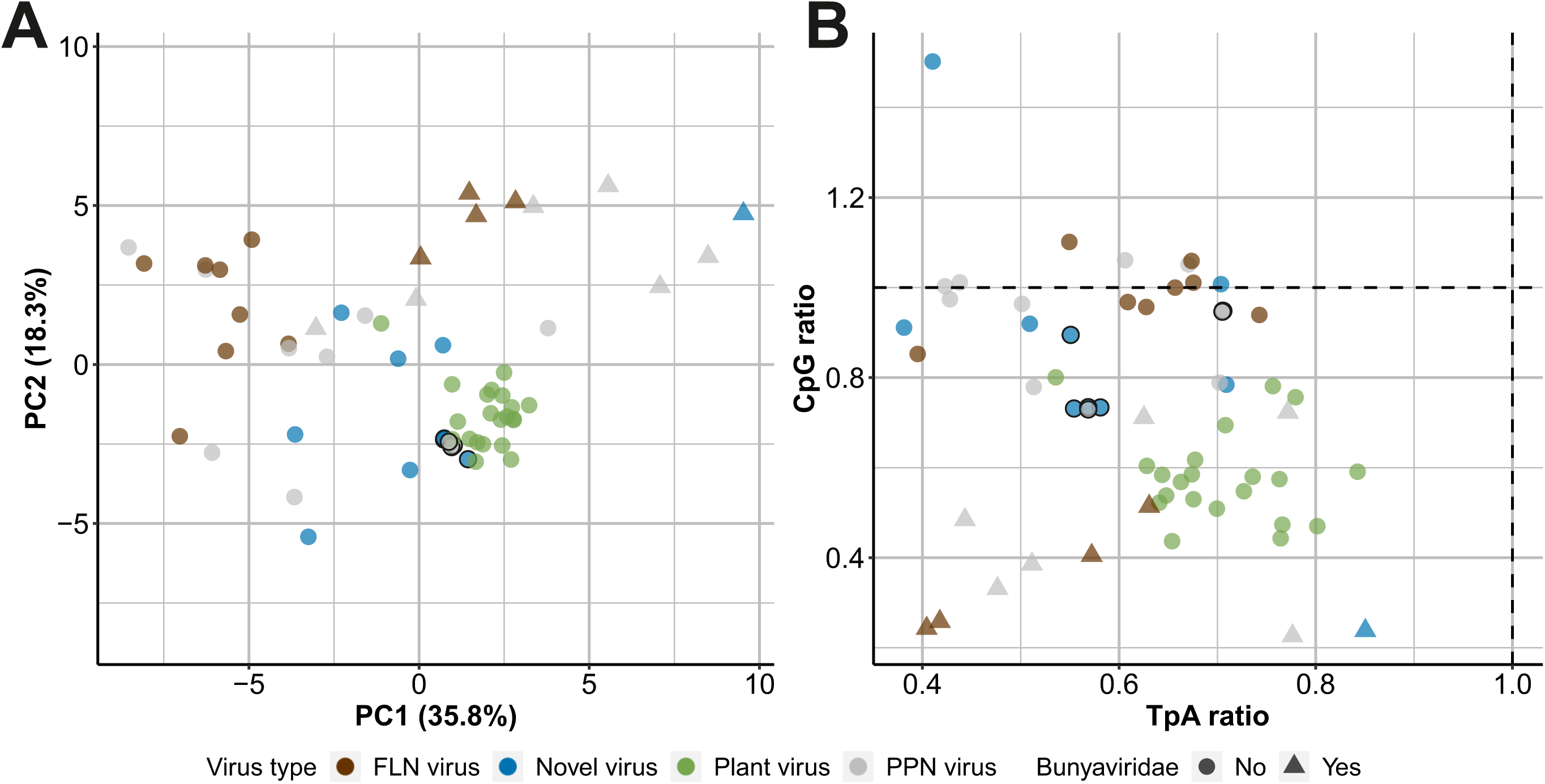
Genome composition analysis of nematode- and plant-infecting viruses. Both plots show information about genomes of viruses infecting free-living nematodes (FLN virus; brown), plant-parasitic nematodes (PPN; grey), plant viruses (Plant virus; green) that are vectored by nematodes and the novel viruses described in this study (Novel virus; blue). See Supplementary Table S3 and S4 for the information about the FLN, PPN and Plant viruses. Viruses with a black outline cluster close to the plant-infecting samples in the PCA and are indicated with a black outline in both plots. Vertebrate infecting viruses belonging to the *Bunyavirales* family are triangle-shaped, all others are circle-shaped. A) Principal Component Analysis (PCA) plot based on the genome composition. The first and second principal component axis (PC1 and PC2) are depicted which together capture 54.1% of the variation in the dataset. B) CpG and UpA observed-to-expected ratios of the viruses. The dashed lines indicate the expected CpG and UpA ratio based on the nucleotide frequency of each genome in the analysis. Samples deviating from these lines have different CpG and UpA ratios than expected indicating potential adaptation to the host species.

Next, CpG_O/E_ and UpA_O/E_ ratios of the same viruses were analysed (**Figure 3B**). The plant-infecting viruses and the vertebrate-infecting viruses from the *Bunyavirales* had lower CpG_O/E_ ratios (≈ 0.6) than the nematode-infecting viruses and most of the new viruses (CpG_O/E_ ratios (≈ 1). Plant- and nematode-infecting viruses are more similar for UpA_O/E_ than CpG_O/E_ ratios. Finally, in this analysis, four viruses that clustered more closely to plant viruses in the PCA, namely the picornaviruses (PCN PLV, SBCNV1) and the Nyami-like virus BCN NLV1, appear more distinct from the plant viruses. Altogether, these analyses, together with the RT-PCR data, suggest that plant- and nematode-infecting viruses can be distinguished based on genome composition analyses and strengthen their identity as genuine nematode-infecting viruses.

### Immune response towards viruses

Based on our analysis of genome composition and phylogenetics, we found that most viruses we studied closely resemble and group with other known nematode-infecting viruses. Our RT-PCR data confirmed the presence of BCN viruses in various BCN Woensdrecht life stages and revealed that PCN viruses were passed from parent to offspring, except for PCN PLV in Ro1-line 19. Moreover, we did not find evidence that any of these viruses were present in the respective genomes of the nematodes. This led us to hypothesize that if these nematode-infecting viruses are actively infecting their hosts, the hosts would likely develop an immune response against them, possibly using RNAi. Therefore, the BCN and PCN nematode’s RNAi-based antiviral immune response was explored by small RNA (sRNA) sequencing of nematode populations. We analysed sRNA profiles of hatched preparasitic J2s of BCN Woensdrecht and PCN Ro5-line 22 and Ro1-line 19 for the presence and frequency of virus-derived small RNAs (viRNA), their first nucleotide bias and their genome coverage.

For BCN Woensdrecht, the viRNA reads represented ca. 0.55% (138269) of the total sRNA reads ranging from 15 to 35nt in length of which 20-24nt fragments indicate products formed by active RNAi of nematodes (Ashe et al., 2013; Coffman et al., 2017; Richaud et al., 2019). From the five viruses infecting BCN Woensdrecht, most viRNAs targeted BCN BLV, while BCN NLV1 and BCN NLV2 were the least targeted by the nematode’s immune system (**Figure 4**). The viRNA profiles of BCN BLV, SBCNV1 and BCN TbLV showed a strong accumulation of 22nt and 23nt reads mapping to the reverse strand of the viral genomes, while the same was not observed for BCN NLV1 and BCN NLV2. Interestingly, irrespective of the virus and the abundance of viRNAs observed, reads ranging from 20 to 24nt in length and longer were strongly biased in their first nucleotide preference towards a G. For the reads shorter than 20nt in length, this bias was less noticeable, and these may represent viral genome degradation products. The predominance of reads with 22nt or 23nt in length resembles the viRNA pattern observed in *C. elegans* against Orsay virus [20,22], albeit for BCN BLV and BCN TbLV the 23nt viRNA accumulated more than the 22nt class, which differs from *C. elegans* infected by Orsay virus where secondary 22nt viRNAs are most prominent [22]. Importantly, we have observed for all viruses that the reads were evenly spread throughout the viral genomes, including for BCN NLV1 and BCN NLV2 with genome coverages ranging from ca. 70% (BCN NLV2) to 95% (SBCNV1) of the respective genome lengths.

**Figure 4.**
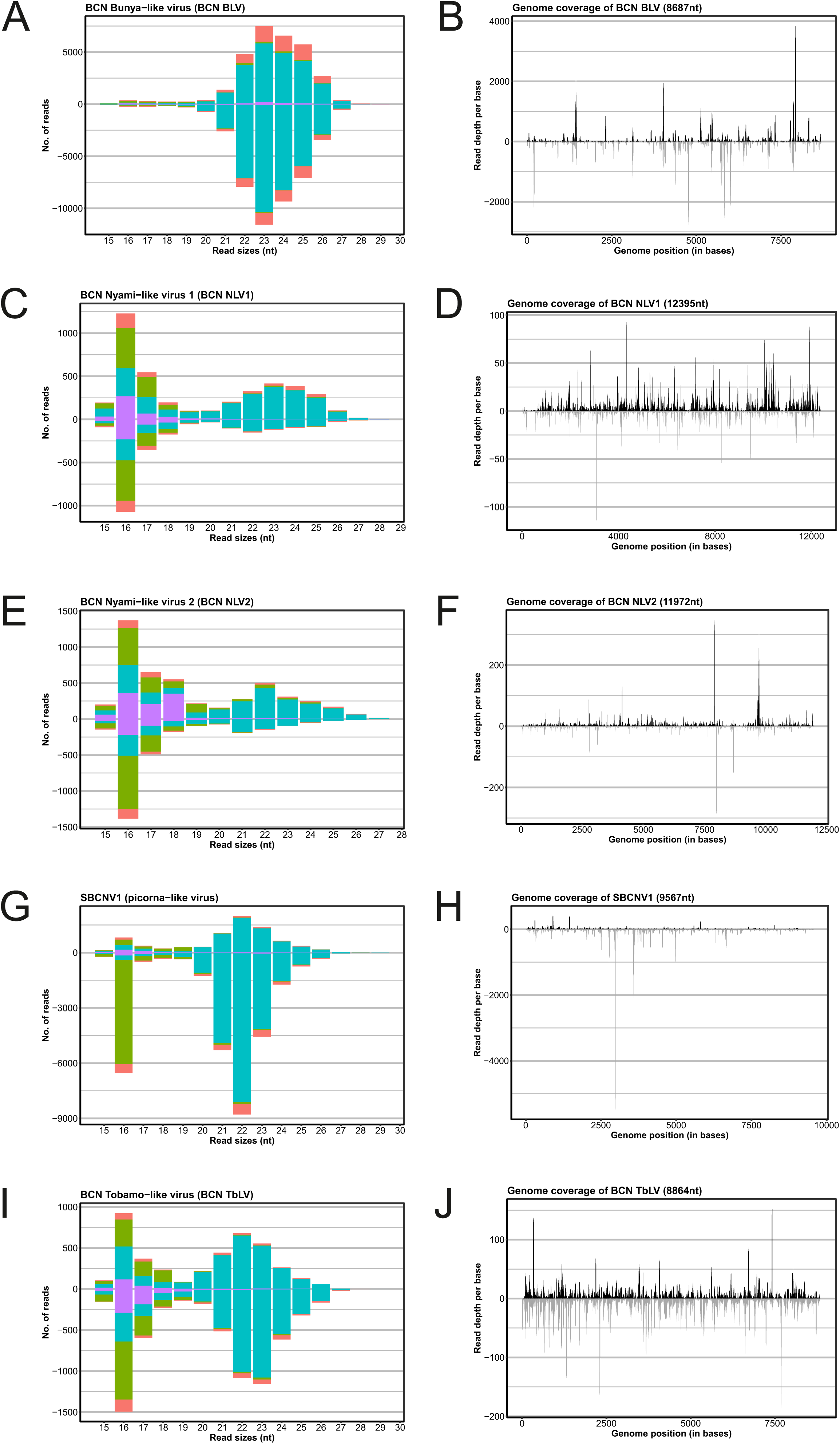
Virus associated small interfering RNA (viRNA) reads of BCN. Read frequency, mapping and first nucleotide bias for each of the viruses identified in beet cyst nematode (BCN Woensdrecht) were determined. Small RNA size distributions contain the total number of reads mapping to each viRNA size category for each virus and indicate the first nucleotide of the mapped sRNA (blue: G, purple: T, green: C, pink: A). The forward mapping reads have positive numbers in the y-axis while the negative mapping reads have negative numbers in the y-axis. The profiles show the genome-wide coverage of viRNA reads throughout the length of each viral genome. **A**) Small RNA size distribution of BCN Bunya-like virus **B**) The profile of BCN Bunya-like virus. **C**) Small RNA size distribution of BCN Nyami-like virus 1. **D**) The profile of BCN Nyami-like virus 1. **E**) Small RNA size distribution of BCN Nyami-like virus 2. **F**) The profile of BCN Nyami-like virus 2. **G**) Small RNA size distribution of SBCNV1. **H**) The profile of SBCNV1. **I**) Small RNA size distribution of BCN Tobamo-like virus. **J**) The profile of BCN Tobamo-like virus.

For PCN the viRNA profiles varied greatly based on the genotype analysed (Ro5-line 22 or Ro1-line 19) with considerably more viRNA reads mapping to the three PCN viruses in *G. rostochiensis* Ro5-line 22 (527710 reads; 2.09% of all sRNA reads generated) than *G. rostochiensis* Ro1-line 19 (11720 reads; ca. 0.05% of all sRNA reads) representing a 41.8 fold difference in viRNA production (**Figure 5, Supplementary Figure S2)**. This large difference in quantities of virus-mapped reads between Ro5-line 22 and Ro1-line 19 reflects the differences in the number of virus-mapped reads in our initial virus discovery between these two lines of PCN (**Table 1**). For PCN Ro5-line 22 the 22nt viRNAs were the most frequent sRNA category, followed closely by 21nt viRNAs and 23nt viRNAs, respectively (**Figure 5**). For PCN PLV and PCN TtLV, the reverse reads were more abundant while for PCN XLV reads mapped slightly more on the positive than on the negative strand. As observed in the analysis of BCN Woensdrecht viRNAs, the reads with 20-24nt in length or longer were heavily biased towards having a G as the starting nucleotide. Viral genome coverages ranged from 84% (PCN PLV) to 100% (PCN TtLV) of the respective bases in the genome. For *G. rostochiensis* Ro1-line 19, no obvious pattern of RNAi response against any of the viruses was observed (**Supplementary Figure S2**). Within this line, a clear bias on the first nucleotide composition of the sRNAs was only observed for PCN TtLV (**Supplementary Figure S2)**. The overall difference in sRNA reads mapping to any of the viral genomes in comparison to Ro5-line 22 is indicative of significant differences in triggering of RNAi in the former line. Nonetheless, the viRNA read coverage varied between 58.8% (PCN PLV) and 90% (PCN TtLV) in Ro1-line 19.

**Figure 5.**
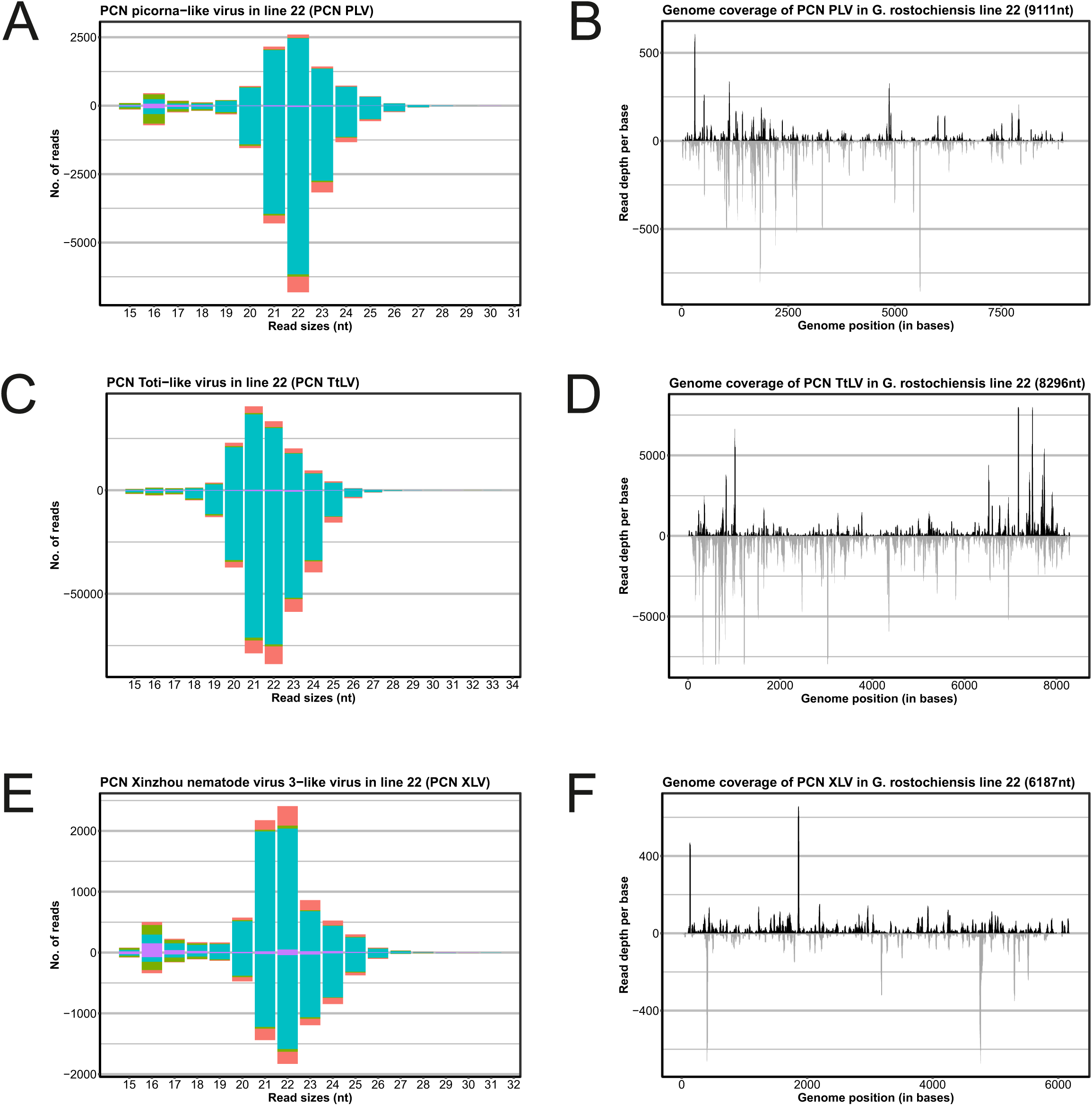
Virus associated small interfering RNA (viRNA) reads of PCN Ro5-line 22. Read frequency, mapping and first nucleotide bias for each of the viruses identified in potato cyst nematode (PCN Ro5-line 22) were determined. Small RNA size distributions contain the total number of reads mapping to each viRNA size category for each virus and indicate the first nucleotide of the mapped sRNA (blue: G, purple: T, green: C, pink: A). The forward mapping reads have positive numbers in the y-axis while the complementary reads (negative) mapping reads have negative numbers in the y-axis. The profiles show the genome-wide coverage of viRNA reads throughout the length of each viral genome. **A**) Small RNA size distribution of PCN Picorna-like virus **B**) The profile of PCN Picorna-like virus. **C**) Small RNA size distribution of PCN Toti-like virus. **D**) The profile of PCN Toti-like virus. **E**) Small RNA size distribution of PCN Xinzhou nematode virus 3-like virus. **F**) The profile of PCN Xinzhou nematode virus 3-like virus.

Taken together, our sRNA analysis indicates that BCN Woensdrecht and PCN *G. rostochiensis* Ro5-line 22 respond to most of the viruses infecting them by producing hallmark viRNAs, albeit in a slightly different fashion than *Caenorhabditis* species [22]. In *G. rostochiensis* Ro1-line 19 no clear pattern of viRNA accumulation was observed, in line with virus-mapping and RT-PCR experiments indicating viruses were less present in Ro1-line 19 than Ro5-line 22. In *G. rostochiensis* Ro5-line 22 PCN TtLV, a double stranded RNA (dsRNA) virus, triggered significantly more viRNAs than any other virus infecting either PCN or BCN. In general, the RNAi response against positive single stranded RNA (+ssRNA) viruses more closely resembled the patterns observed in *C. elegans* (against the +ssRNA Orsay virus) than the RNAi responses against negative single stranded (-ssRNA) viruses.

We observed a striking differences in viRNA profiles and viRNA presence/absence in the two investigated PCN lines. Furthermore the RT-PCR data that indicated that PCN PLV and PCN XLV were present in Ro1-line 19 confirmed this observation. Therefore, we investigated the presence of orthologs of genes involved in the antiviral immunity in *C. elegans* between Ro1-line 19 and Ro5-line 22 (**Supplementary Table S5**). For eight genes, orthologues could be mapped in *G. rostochiensis*. Of these, orthologues to *dcr-1*, *nck-1*, and *rde-8* were present in both Ro1-line 19 and Ro5-line 22. Four orthologues had distinct orthologue numbers in either Ro1-line 19 and Ro5-line 22: RNAi genes *drh-1* (one additional orthologue in Ro1-line 19), *rrf-3* (one additional distant related orthologue in Ro5-line 22) and *eri-1* (two additional orthologues in Ro5-line 22), and the STAT pathway gene *sta-1* (two additional orthologues in Ro-1 line 19). Finally, for the kinase *sid-3* an orthologue was only found in Ro5-line 22, but not in Ro1-line 19 (**Supplementary Figure S3**). Presence of *sid-3* makes *C. elegans* 100-fold more permissive to viral infection and is required for viral entry [60,61]. If *sid-3* has a similar function in cyst nematodes, absence of a *sid-3* orthologue in Ro1-line 19 might explain why this line is less susceptible to viral infection.

## Discussion

### PPN mount an active RNAi response towards viruses

This research explored the presence of viruses and an active antiviral response in in two cyst nematode species: *Heterodera schachtii* (BCN Woensdrecht) and *Globodera rostochiensis* (PCN Ro1-line 19 and Ro5-line 22). To establish whether the discovered viruses infect PPN rather than being cryptic or “hitchhiking”, we investigated the antiviral RNAi-response. We found distinct patterns of RNAi response in BCN and PCN against viruses of positive or negative strand orientation: the immune response against -ssRNA viruses did not show a strong bias towards the 22G hallmark viRNAs contrary to the +ssRNA viruses we investigated. This could indicate that either the -ssRNA viruses are not targeted by RNAi, or that they did not replicate efficiently in these nematodes. Moreover, we observed genotype-specific differences in viral infection between two *G. rostochiensis* lines that might relate to genetic diversity in the RNAi pathway genes.

Our analysis of the RNAi responses against viruses is the first of its kind for any PPN virus and it indicates that viRNA patterns of immune responses observed in *Caenorhabditis* also occur in cyst nematodes against some, but not all their infecting viruses. Our data indicated that BCN Woensdrecht reacted to +ssRNA viruses and -ssRNA viruses differently in terms of the presence and abundance of characteristic 22G viRNAs with clear patterns of RNAi response observed against the +ssRNA viruses, but not against the -ssRNA viruses. A similar observation, regarding the abundance of the 22G viRNAs, was made for the small RNA pattern generated in PCN Ro5-line 22 against the -ssRNA virus PCN XLV yielding relatively few viRNAs without strong antisense bias. These analyses suggest differences may exist in the RNAi responses of BCN and PCN against +ssRNA, (dsRNA) and -ssRNA viruses, the latter containing fewer or none of the hallmarks that resemble those of a classical antiviral RNAi immune response in nematodes [20–22]. These findings shed light on the function of the antiviral response in plant-parasitic nematodes against specific viruses as well as the interplay between nematode immune responses and potential silencing suppression or immune evasion by their viral counterparts.

### Potential interactions between viruses and the molecular RNAi machinery in PPN

While no evidence exists indicating that any of the viral proteins in Nyami-like viruses or PCN XLV or the L protein of BCN BLV have silencing suppression activity, these types of mechanisms were found for several members of the *Bunyavirales* order [62]. Silencing suppressor proteins might disrupt the host RNAi response by i) acting specifically to prevent the loading of small RNAs into RISC complexes or ii) inhibiting the amplification of the RNAi response by interfering or competing with the host RDRs for viral RNA substrates [62]. On the other hand, the genetic makeup of BCN might be involved in the differences observed in small RNA patterns. It is also possible in the case of the -ssRNA Nyami-like viruses BCN NLV1 and BCN NLV2 that these viruses replicate in the nucleus and therefore trigger different intracellular RNAi responses than viruses that are cytoplasm-bound [63]. This may generate different patterns of small interfering RNA accumulation, alike RNAi patterns observed in plants. PCN PLV and PCN XLV appear to accumulate to lower levels in Ro1-line 19 than in Ro5-line 22. According to our comparative genomic analysis, the difference in viral susceptibility observed between these two lines may be accounted for by the absence of *sid-3* in Ro1-line 19 as this gene enhances susceptibility to viral infection in *C. elegans* [60,61]. Importantly, absence of *sid-3* should still be confirmed experimentally as current gene predictions may be incorrect. Nevertheless, the lack of an orthologue of *sid-3* might explain the stark contrast in viral accumulation and in viRNA profiles. Thus, both the type of virus as well as the genetic makeup of the nematode species appear to determine the RNAi response.

### Plant-parasitic nematodes carry multiple viruses

Both PCN and BCN were coinfected with respectively three and five viruses, with BCN carry two distinct species of Nyami-like viruses. Coinfection of nematodes with multiple viruses was previously described for *C. briggsae* [64] that can be infected with Le Blanc and Santeuil virus simultaneously. These nodaviruses are even capable to infect a single cell within an individual. Whether Le Blanc and Santeuil virus, and different strains of Santeuil, competed within the host depended on the host genotype [64]. The host genotype determines its RNAi capacity and we hypothesize that this may determine outcomes of coinfection in nematodes as mosquito cells were better protected against superinfection (infection by a second virus) when they lacked a potent RNAi response [65]. Additionally, superinfection of *C. elegans* with two variants of the Orsay virus indicates that secondary infections produced less viRNAs than acute primary infections [66]. Future (super)infection experiments, in particular using *G. rostochiensis* Ro1-line 19 and Ro5-line 22 that showed distinct viral profiles and RNAi responses, could reveal within-host interactions between the viruses present.

### Fitness costs and transmission mechanisms of nematode-infecting viruses

Viruses could potentially be applied as biocontrol measures against PPN, and it is therefore important to assess their impact on host fitness and their modes of transmission. Currently there is no indication in literature of fitness costs or benefits for the nematode associated to viral infection in PPN. Our findings that RNAi is triggered against these viruses are the first to point to a potential fitness cost associated to viral parasitism in PPN. For *C. elegans*, Orsay virus infection appears to cause only mild fitness effects [20,67], although some nematode lines slow down growth [64]. Other experiments conducted on Orsay viruses indicate that naïve males preferentially mated with uninfected hermaphrodites in comparison to virus-infected hermaphrodites [68]. Given that we find infections in young adult and older adult females of BCN Woensdrecht for the five viruses identified in this population, we aim to conduct further research exploring potential effects on mating preferences and/or fecundity of infected females and naïve females and vice-versa. Based on the biology and life cycle of sedentary endoparasitic nematodes, the possible routes for virus acquisition are via their plant hosts, horizontal transmission within the cyst during hatching and/or J2 movement towards the host plant roots, and finally sexual (vertical or horizontal) transmission during fecundation.

Mounting evidence from different viruses infecting *Caenorhabditis* nematodes indicates that nematode viruses may be horizontally (Orsay, Le Blanc viruses) or vertically transmitted (Bunya-like viruses in *C. remannei*, *C. briggsae* and *C. zanzibari*) [11,21,64,69]. Moreover, the accumulation of a -ssRNA rhabdovirus in the ovaries and testes of the human-parasitic nematode *Onchocerca volvulus* female and male adults indicated that this virus is sexually transmitted [53]. Interestingly, the Bunya-like *Caenorhabditis* viruses [21] and the BCN BLV described here latter differ from most viruses in the order *Bunyavirales*, as they are missing their M and S RNA segments which might be an indication of an adaption towards a different mode of transmission. The remarkably high conservation between the SCBNV1 variant found in BCN populations grown in Tennessee and our ‘Woensdrecht’ population could suggest a common origin [10]. Our analysis of *G. rostochiensis* Ro5-line 22 and Ro1-line 19 suggests that viruses in PPN may be horizontally or sexually transmitted. PCN PLV, PCN XLV, and PCN TtLV are all present in Ro5-line 22 and in its ancestral population (PCN Harmerz) while for Ro1-line 19, out of the two viruses confirmed to be present (PCN PLV and PCN XLV) only one (PCN XLV) is found in its ancestral population (Mierenbos). These findings indicate that PCN XLV may be sexually or vertically transmitted. Conversely, PCN PLV may (also) be horizontally transmitted across individuals as it was independently acquired by Ro1-line 19 during cyst multiplication.

## Conclusions

There still exist many unanswered questions about the effects that PPN viruses have on their host, most critically about the associated fitness costs. Here we have for the first time described an antiviral RNAi response in two PPN species, which is indicative of active viral infections and possible fitness costs. We have also gathered indications that viruses might be acquired both sexually and horizontally. Future research should aim at generating and understanding differences between virus-infected and virus-free (or cured) nematode populations. Of particular interest is the difference in viral accumulation between *G. rostochiensis* Ro1-line 19 and Ro5-line 22. Follow-up genomic studies would benefit from using these two lines, because of the extensive previous knowledge existing on the development and genetics of these inbred lines. In this way, the functioning of the immune system of PPN can be further elucidated, which is a key element in understanding PPN biology and may lead to new research directions in PPN control.

---

**Supplemental Figure 1 RdRp or polyprotein based phylogenetic analyses of novel BCN (A to C) and PCN (D to F) viruses. –** All trees contain the same indications: viral taxa that are nematode-associated are highlighted by a colour strip on the right side of each tree in purple. Within the nematode-associated taxa, a second colour scheme is applied to differentiate between nematode-associated viruses. Highlighted in light blue and in bold letter are the novel viruses described in this manuscript. Previously known PPN viruses are highlighted in grey. Animal parasitic nematode-associated viruses are highlighted in pink and free-living nematode-associated viruses are highlighted in yellow or brown. For the phylogenetic analysis a minimum bootstrap support level was set at 70. Branches below this value were removed from the tree.

**Supplementary Figure 2 Virus associated small interfering RNA (viRNA) reads of PCN Ro1-line 19 –** Read frequency, mapping and first nucleotide bias for each of the viruses identified in potato cyst nematode (PCN Ro1-line 19) were determined. Small RNA size distributions contain the total number of reads mapping to each vsiRNA size category for each virus and indicate the first nucleotide of the mapped sRNA (blue: G, purple: T, green: C, pink: A). The forward mapping reads have positive numbers in the y-axis while the complementary reads (negative) mapping reads have negative numbers in the y-axis. The profiles show the genome-wide coverage of viRNA reads throughout the length of each viral genome. **A**) Small RNA size distribution of PCN Picorna-like virus **B**) The profile of PCN Picorna-like virus. **C**) Small RNA size distribution of PCN Toti-like virus. **D**) The profile of PCN Toti-like virus. **E**) Small RNA size distribution of PCN Xinzhao virus 3-like virus. **F**) The profile of PCN Xinzhao virus 3-like virus.

**Supplementary Figure 3 A phylogenetic tree of *sid-3* in different nematode species –** Orthologs of *C. elegans sid-3* were found in diverse nematode populations, including *H. schachtii* populations and *G. rostochiensis* Ro5-line 22, but excluding *G. rostochiensis* Ro1-line 19.

## Acknowledgements

The authors thank Richard Kormelink and José E. Pacheco Diaz for providing feedback on the manuscript. Dennie te Molder is thanked for his advice for the PCA analysis and Jelke Fros for his help with CpG analyses. Lisa van Sluijs was supported by Sectorplan BÈTA-II Biology. Jaap-Jan Willig was supported by Dutch Top Sector Horticulture & Starting Materials (TU18152). André Bertran and Hein Overmars were supported by NWO domain Applied and Engineering Sciences H.I.P. grant BB1.1 (16873). M.G.S. was supported by NWO domain Applied and Engineering Sciences VENI grant (17282) and VIDI grant (21240).

## Credit author statement

*André Bertran*: Conceptualization, Methodology, Software, Validation, Formal analysis, Investigation, Resources, Data curation, Writing – original draft, review and editing, Visualization, Supervision, Project administration *Lisa van Sluijs*: Conceptualization, Methodology, Software, Validation, Formal analysis, Investigation, Resources, Data curation, Writing – original draft, review and editing, Visualization, Project administration, Funding acquisition *Hein Overmars*: Conceptualization, Methodology, Validation, Formal analysis, Investigation, Resources, Writing – original draft, review and editing, Visualization, Supervision *Matthijs Mobach*: Formal analysis, Investigation, Resources *Stefan van de Ruitenbeek*: Methodology, Software, Validation, Formal analysis, Investigation, Resources, Data curation, Writing – original draft *Jaap-Jan Willig*: Resources *Joris van Steenbrugge*: Methodology, Software, Formal analysis, Resources *Aska Goverse*: Conceptualization, Resources, Writing – Review and Editing, Supervision, Project administration, Funding acquisition *Mark Sterken*: Conceptualization, Resources, Writing – Review and Editing, Supervision, Project administration, Funding acquisition

