## Supplementary figures and images for "Plant parasitic cyst nematodes respond to viral infection through RNA interference"

### Supplementary Figure S1

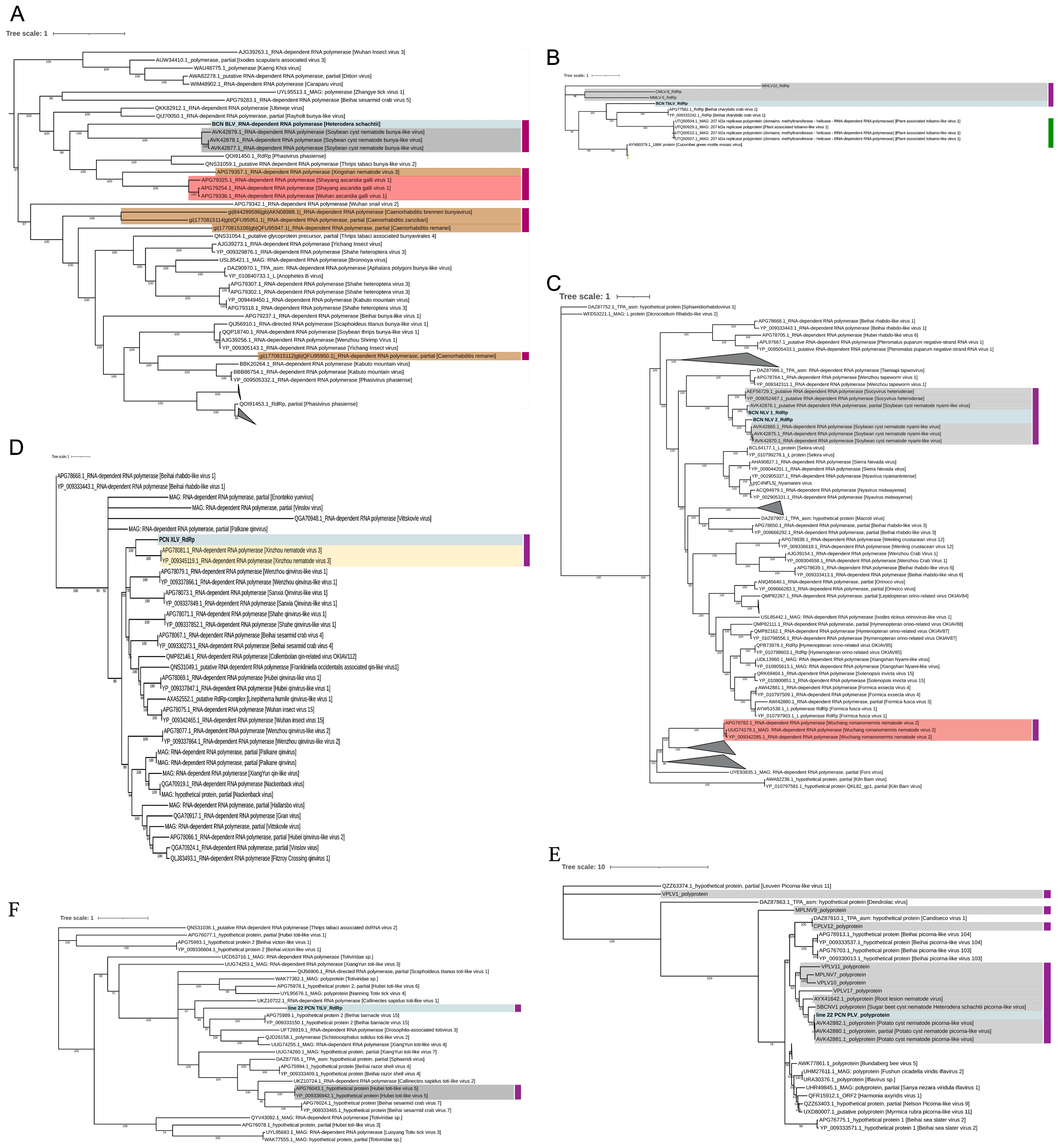

### Supplementary Figure S2

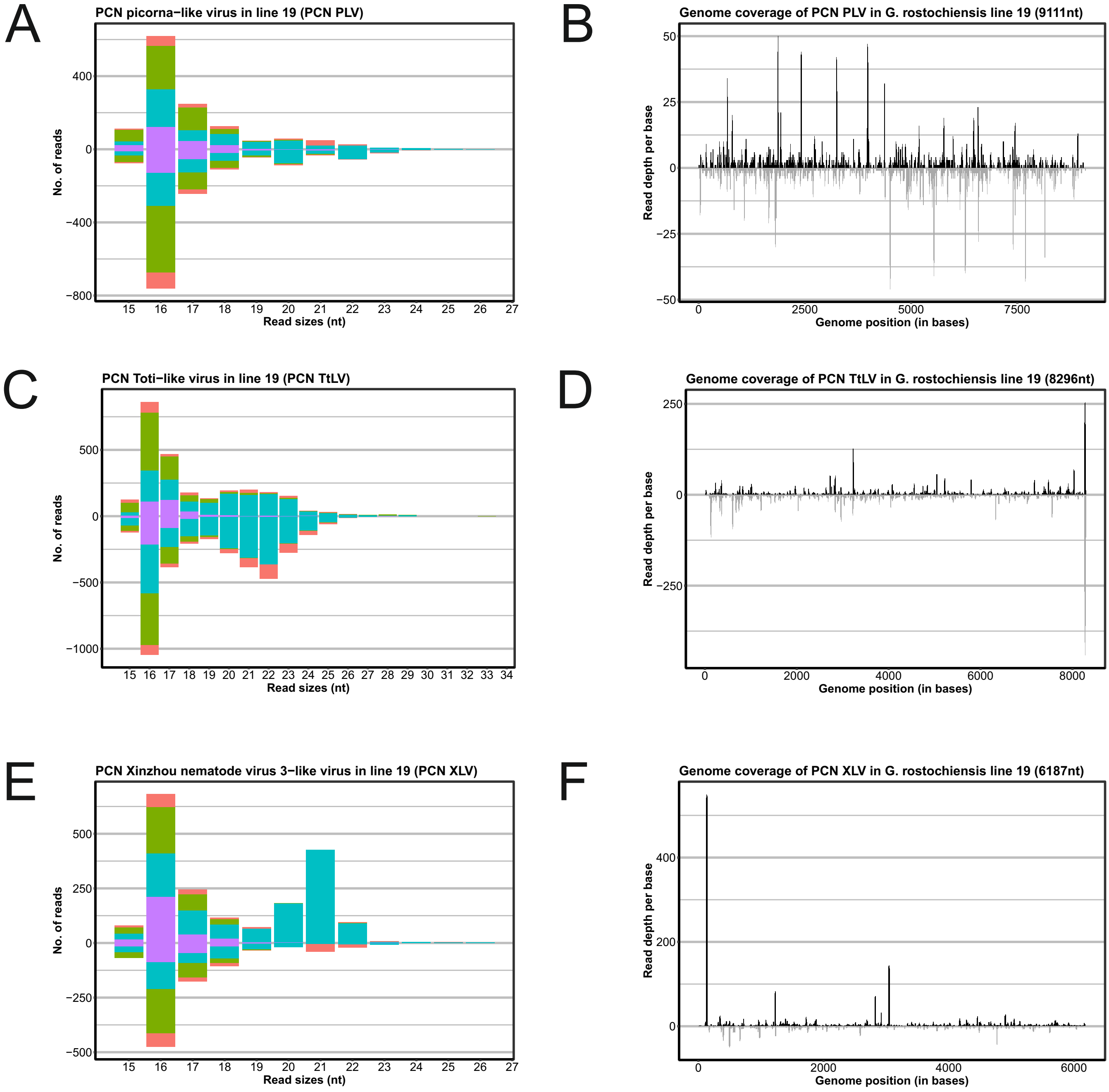

### Supplementary Figure S3

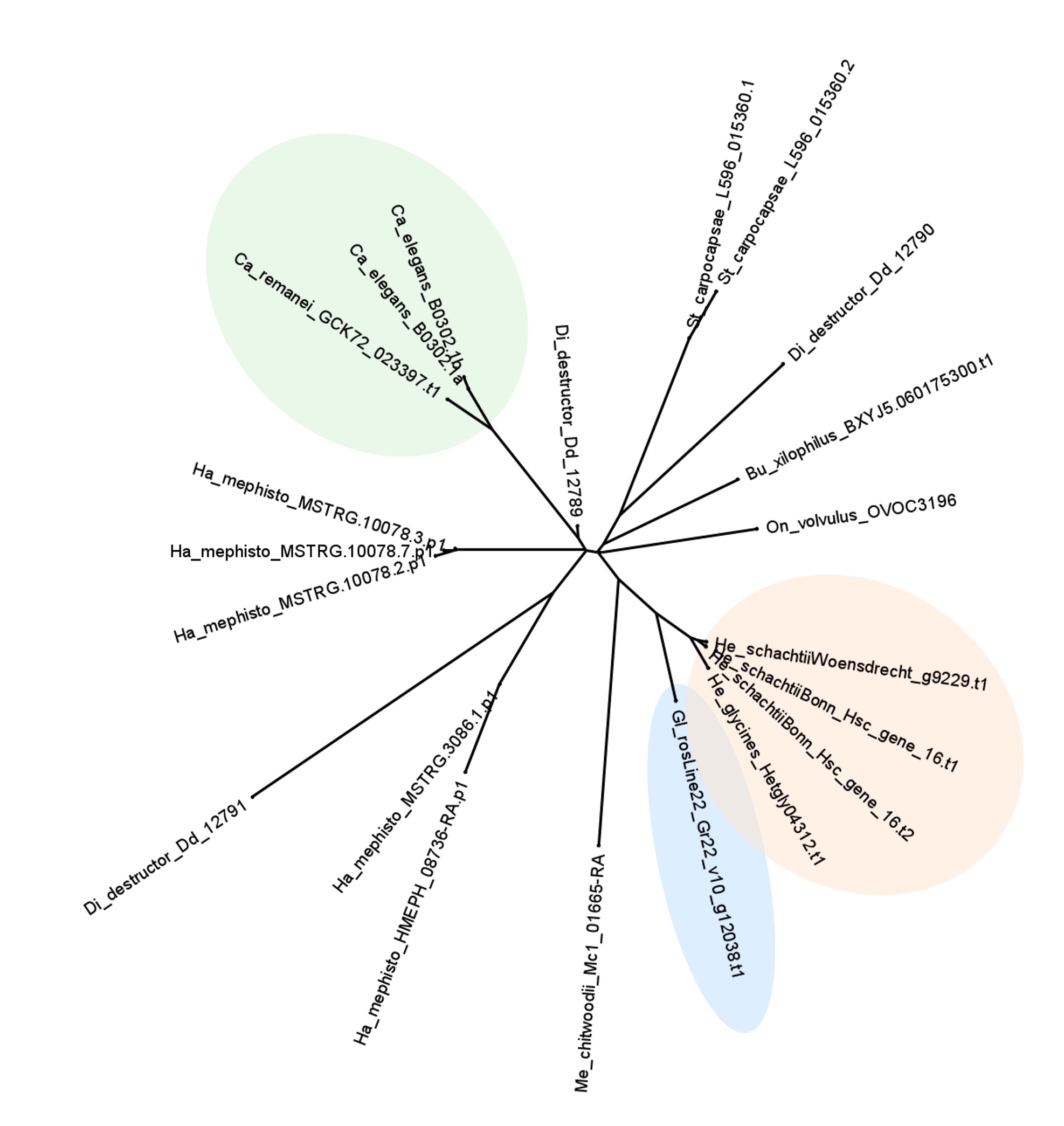
